# Stratifying risk of prostate cancer recurrence following external beam radiation therapy: Comparing prostate MRI with prostate biopsy pathology

**DOI:** 10.1101/056879

**Authors:** William Millard, Jason Slater, Samuel Randolph, Thomas Kelly, David Bush

**Affiliations:** Loma Linda University, Department of Radiology.; Loma Linda University, Radiation Oncology.

**Keywords:** Percent cancer volume, percentage of positive biopsy cores, dominant intraprostatic lesion, prostatic neoplasms/pathology, external beam radiation therapy

## Abbreviations

(DIPL): Dominant intraprostatic lesion
(DN): Dominant nodule
(EBRT): External Beam Radiation Therapy
(ECE): Extracapsular Extension
(NVB): Neurovascular Bundle Invasion
(SVI): Seminal Vesicle Invasion
(MIBC): Maximum involvement of any biopsy core
(PCV): Percentage of cancer volume
(PPBC): Percentage of positive biopsy cores
(NCCN): National Comprehensive Cancer Network

## INTRODUCTION

Prostate cancer and therapy for the disease is an important cause of morbidity and mortality in the developed world. Reducing over-treatment and under-treatment are areas of particular interest in the modern approach to prostate cancer. Typical therapeutic approaches include active surveillance, radical prostatectomy, and radiation therapy, with hormonal therapy playing an adjunctive role. Treatment strategies for prostate cancer are often informed by appropriate risk stratification of the patient. Multiple risk stratification tools have been developed including that of D’Amico and colleagues which separates patients into low, intermediate, and high risk categories based on pre-treatment PSA, prostate core biopsy Gleason score, and clinical T-stagingd^1^. This has formed the basis of many of the newer stratification tools such as the National Comprehensive Cancer Network (NCCN) classification, which also includes a very low risk category^2^. There have been several attempts recently to refine the risk stratification systems to more accurately identify patients at lower and higher risk. These methods include additional clinical information such as information from prostate biopsy cores as well as prostate MR findings used to further subdivide risk and predict treatment outcomes.^3–8^

Recent work at our institution comparing several novel prostate core biopsy findings in low and intermediate risk patients identified a subset of clinically localized prostate cancer patients in the intermediate risk category who were more likely to fail external beam radiation therapy.^4^ Some patients in this category respond more like high risk patients with regard to post EBRT outcomes. Identifying these cases at the onset of therapy could lead to improved meaningful outcomes. Additionally, research using findings on pre-treatment prostate MRI has demonstrated these to independently predict post-EBRT PSA relapse.^9^ Comparing MRI and pathological data may help to accurately further risk stratify patients undergoing EBRT. Our purpose here is to determine if there is a correlation between the poor prognostic factors demonstrated on prostate biopsy cores and selected findings on prostate MRI.

## MATERIALS AND METHODS

A group of patients with clinically localized intermediate risk category prostate cancer with 1.5 T and 3.0 T pretreatment MRI scans performed during a predefined time frame (2007–2011) were selected for a retrospective cohort study. Approval was obtained for this study from the Loma Linda University Institutional Review Board. 65 cases were reviewed by 2 body trained radiologists with 22 years and 5 years of experience. Readers were aware the patients had prostate cancer as per routine, but were blinded to other clinical information such as risk stratification or patient outcomes. Reader consensus was obtained at the time of readout regarding presence of extracapsular extension, seminal vesicle invasion, and presence of disease in each sextant by T2 and ADC imaging, as well as presence of one or more sextants containing an intraprostatic dominant nodule. Results were analyzed for correlation between these findings on MRI and the results of pre-treatment prostate biopsy cores which included percentage of cancer volume (PCV), percentage of positive biopsy cores (PPBC), and maximum involvement of any biopsy core (MIBC). Statistical analysis was performed with Pearson correlations and independent samples t-test with p-values less than 0.05 considered significant. Analysis was performed using SPSS statistical software.

## RESULTS

The average age of participant in this study was 66.5 years (range 48.0–81.0). The cohort consisted entirely of intermediate risk patients per the NCCN risk categorization.^2^ Other relevant demographic data is listed in Table 1. Of the 65 patients, all but one patient were found to have T2 hypointense foci in at least one sextant of the prostate. One or more sextants containing a dominant nodule were identified in 40 of 65 patients. Extracapsular extension, neurovascular bundle invasion, and seminal vesicle involvement were present in 10, 11, and 21 patients respectively. This is summarized in Table 2.

**Table 1.** Demographics.

|  | Count/Mean (Range) |
| --- | --- |
| Total Patients | 65 |
| Age | 66.5 (48.0-81.0) |
| Pre-EBRT PSA | 7.25 (0.70-16.5) |
| T Stage |  |
| T1c | 34 |
| T2a | 16 |
| T2b | 11 |
| T2c | 4 |
| Gleason Score |  |
| 3+3=6 | 9 |
| 3+4=7 | 41 |
| 4+3=7 | 15 |

**Table 2.** MR Findings.

|  | Count |
| --- | --- |
| T2 Hypointense Foci |  |
| 0 | 1 |
| 1 | 2 |
| 2 | 3 |
| 3 | 3 |
| 4 | 11 |
| 5 | 9 |
| 6 | 36 |
| Dominant Foci |  |
| 0 | 25 |
| 1 | 17 |
| 2 | 19 |
| 3 | 4 |
| Extracapsular Extension |  |
| Present | 10 |
| Absent | 55 |
| Neurovascular Bundle Invasion |  |
| Present | 11 |
| Absent | 54 |
| Seminal Vesical Invasion |  |
| Present | 21 |
| Absent | 44 |

PPBC was statistically significantly positively correlated with greater number of T2 hypointense prostate sextants, greater number of sextants containing a dominant nodule, and absolute presence of a dominant nodule. PCV similarly positively correlated with greater number of sextants with a dominant nodule and absolute presence of a dominant nodule, however PCV was negatively correlated with seminal vesicle invasion and combined presence of advanced disease (ECE, NVB, or SVI). MIBC was positively correlated with greater number of sextants with a dominant nodule, but no other finding. These correlations are summarized in Table 3.

**Table 3.** Pearson correlations between MRI findings and Prostate Biopsy Cores.

|  |  | # of T2 foci | # of DN | ECE Y/N | NVB Y/N | SVI Y/N | Advanced Disease Y/N | DN Y/N |
| --- | --- | --- | --- | --- | --- | --- | --- | --- |
| PPBC | Pearson Corr | 0.27 | 0.53 | -0.01 | 0.02 | -0.2 | -0.17 | 0.33 |
|  | Sig. (2-tailed) | 0.029 | <0.005 | 0.907 | 0.896 | 0.119 | 0.175 | 0.006 |
| PCV | Pearson Corr | 0.19 | 0.47 | -0.08 | -0.08 | -0.26 | -0.33 | 0.29 |
|  | Sig. (2-tailed) | 0.135 | <0.005 | 0.525 | 0.528 | 0.034 | 0.008 | 0.019 |
| MIBC | Pearson Corr | 0.04 | 0.27 | -0.03 | 0 | -0.05 | -0.16 | 0.22 |
|  | Sig. (2-tailed) | 0.733 | 0.027 | 0.824 | 0.998 | 0.669 | 0.207 | 0.076 |

The presence of a dominant nodule was statistically significantly associated with higher mean PPBC and PCV values (p-values of 0.006 and 0.019 respectively). These had a mean increase of 16% for PPBC and 8% for PCV (Table 4). PSA, Gleason score, and MIBC were not found to be correlated with absolute presence of a dominant nodule on MRI (Not shown).

**Table 4.** Independent samples T test comparing presence of a DN

|  | Presence of a dominant nodule | N | Mean | Std. Deviation | S.E. Mean | Mean Difference | Sig. (2-tailed) |
| --- | --- | --- | --- | --- | --- | --- | --- |
| PPBC | NO | 25 | 0.4 | 0.18 | 0.04 | 0.16 | 0.006 |
|  | YES | 40 | 0.56 | 0.23 | 0.04 |  |  |
| PCV | NO | 25 | 0.15 | 0.12 | 0.02 | 0.08 | 0.019 |
|  | YES | 40 | 0.23 | 0.14 | 0.02 |  |  |
| MIBC | NO | 25 | 0.49 | 0.28 | 0.06 | 0.13 | 0.076 |
|  | YES | 40 | 0.62 | 0.27 | 0.04 |  |  |

## DISCUSSION

NCCN intermediate risk category prostate cancer patients represent a progressively less homogenous group in which the addition of MRI has the potential to play a significant role in further risk stratification. Based on previous research done at our institution, it was found that intermediate risk patients with PPBC >66.7% or PCV >22.5% had higher rates of biochemical failure following external beam radiation therapy. The exact reason for this is not yet known. One hypothesis for the mechanism of these poorer outcomes is that patients with these adverse pathologic risk factors are actually harboring more locally advanced disease that is undetectable on clinical physical exam alone. In such cases, addition of MRI could add greater sensitivity for detection of these abnormalities. Indeed, in our study we found 25 of the total 65 patients (38%) identified as having clinical T1 or T2 stage disease prior to MRI would be categorized as having T3 disease after completing pre-treatment MR imaging. As a result, these patients would more appropriately be classified in the high risk category for prostate cancer. This effect following prostate MRI is not new to the literature. The upstaging of patients has been described previously in work by Riaz et al, where it was found that 61% of their total cohort were classified as having greater T-stage of disease following MRI. This unequally affected those patients in the lower T stage category to a greater degree, with 85% of T1 patients upstaged, 48% of T2 patients, and 7% of T3 patients.^10^ Counterintuitively however, in the cohort we studied those patients with T3 disease had a negative correlation with PCV (Pearson correlation −0.33, p=0.008) and a statistically insignificant negative trend with PPBC. This suggests that the increased MRI T stage is not likely to be the best explanation for increased PPBC and PCV and the resulting increased risk of prostate cancer recurrence previously identified.

Another hypothesis for the relationship between increased risk of biochemical disease recurrence and PPBC and PCV may be related to the morphology of cancer in the prostate. In our study we found that the strongest correlation between the PCV and PPBC was what we termed a dominant nodule. This finding is not inherently new in our study design. Variations of this concept have been described in the literature using terms such as “dominant intraprostatic lesion,” “dominant tumor,” and “index lesion.”^9,11–14^ Much of the interest in this type of MRI prostate finding is found in the radiation oncology literature.^9,11,12^ The exact definition of a dominant nodule is slightly variable but in our study we defined it as a moderately well-defined focus of T2 hypointensity within the prostate peripheral zone. This typically had a gradient of demarcation with adjacent more normal appearing gland. The nodule may involve one or more sextants of the prostate. By contrast, any area of ill-defined T2 hypointensity within a prostate sextant was designated as having probable tumor infiltration without a nodular morphology and was classified as simply T2 hypointense. There was a statistically significant positive correlation between these peripheral zone T2 hypointensities and PPBC (r= 0.27, p=0.029), however the correlation was stronger and included PCV and MIBC for number of sextants containing dominant nodule (PPBC r= 0.53, p=<0.005; PCV r= 0.47, p=<0.005; MIBC r= 0.0.27, p=0.027). In our work, the dominant nodule has been found to have the greatest correlation with the PCV and PPBC of the MRI findings studied. This indicates the dominant nodule may be the underlying mechanism for the higher risk of biological disease recurrence previously identified. If so, the dominant nodule is important as it has potential therapeutic implications. For example, there is developing interest in local dose escalation on Dominant Intraprostatic Lesions (DIPL) which relies on precise localization using MRI.^12^ There are reports of up to 98% 10 year disease free survival rate in patients with DIPL who have undergone a targeted therapeutic approach.^11^ The association of increased PCV and PPBC with a dominant nodule has the potential to help select patients with greater likelihood of having a DIPL for pretreatment MRI. Identification could provide additional prognostic information in these patients and select the best candidates for newer treatment protocols such as focal dose escalation.

Several articles have been published supporting MRI for additional independent risk stratification of patients prior to therapy. In 1996 D’Amico demonstrated that MRI was excellent in identifying extracapsular extension (ECE) and seminal vesicle invasion (SVI) that would otherwise be missed using the more traditional clinical markers of the time.^15^ More recently, Fuchsjäger et al. found several of variables to be statistically significant predictors of outcome in prostate cancer patients receiving EBRT, including MRI TN stage, extracapsular extension (ECE), the number of sextants involved by all lesions, presence of tumor in the apex of the gland, number of sextants involved by the index lesion, the index lesion diameter and the index lesion volume.^9^ The index lesion was described as the “dominant lesion” in their study. While only ECE remained significant in multivariate analysis, the hazard ratio for this finding was notable at 3.04. Similar work by Riaz et al. showed ECE to correlate with outcomes in patients receiving combined EBRT and brachytherapy.^10^ Park, et al, found that among patients who underwent radical prostatectomy, a greater total number of T2, ADC, and DCE features correlated with a larger hazard ratio for biochemical disease recurrence.^16^ Still, challenges remain with regard to widespread use of MRI for additional risk stratification in patients undergoing EBRT. MRI is not widely used in patients with low and intermediate risk disease, and financial constraints make these studies difficult to obtain in some settings. Still, pre-treatment MRI has considerable relevant prognostic and staging information to convey and identifying methods to select patients that will benefit most from pre-treatment MRI would be of considerable benefit.

While our results are very promising, this work is not without its limitations. Firstly, the study uses retrospective design which inherently limits the predictive power of the analysis. However as we are not attempting to predict patient outcomes in this phase of the work, this research design can help direct future work. Additionally, we have taken typical measures to avoid reader bias using reader blinding and consensus between readers. We have a modest number of patients in our initial analysis, and followup work will benefit from a greater number of cases. Re-evaluation of each MRI case is time intensive as each case must be read de novo and consensus obtained, and the process is ongoing. There is the risk of not including statistically significant correlations among other findings we evaluated. Due to the strength of the associations thus far however, we suspect that any statistically significant correlations missed may not be as clinically significant. Probably the greatest limitation is the lack of outcome data available to correlate with. Because of the relatively short time that has elapsed between acquisition of MRI/treatment initiation and the time of the study, there are few treatment failures, thereby preventing a representative cohort of failures and successes to compare. This was not our original intention and does not diminish the results obtained. It is our intention to address this shortcoming in the future.

The strong correlation between the dominant nodule, PCV and PPBC in our study raises the question of whether the presence of a dominant nodule on MRI could be part of the underlying pathophysiology that ultimately results in patients with these adverse findings having higher rates of PSA failure following EBRT for prostate cancer. To our knowledge, there has been little research examining the correlation between prostate biopsy markers and MRI findings. Further work to analyze the relationship between our findings and patient outcome data is ongoing.

## CONCLUSION

We have identified a strong positive correlation between MRI findings of a dominant nodule and prognostic biopsy findings of elevated PPBC and PCV. Further work using these findings in conjunction may aid in further risk stratifying patients at greater risk of post-EBRT PSA relapse.

